# Screening the MMV Open Access Pathogen box unveils novel and potent inhibitors of Amoebiasis agent: *Entamoeba histolytica*

**DOI:** 10.1101/723361

**Authors:** Rufin Marie Kouipou Toghueo, Darline Dize, Benoît Laleu, Patrick Valere Tsouh Fokou, Eugenie Aimee Madiesse Kemgne, Fabrice Fekam Boyom

**Affiliations:** Antimicrobial and Biocontrol Agents Unit (AmBcAU), Laboratory for Phytobiochemistry and Medicinal Plants Studies, Department of Biochemistry, Faculty of Science, University of Yaoundé I, P.O. Box 812, Yaoundé, Cameroon; Medicines for Malaria Venture, Route de Pré-Bois 20, PO Box 1826, 1215 Geneva, Switzerland

**Keywords:** MMV Pathogen Box, Anti-*Entamoeba histolytica*, Novel scaffolds, Lead compounds

## Abstract

Amoebiasis caused by the protozoan parasite *Entamoeba histolytica* remains a major public health hazard, as being the second cause of death among parasitic infections. Although currently prescribed drugs have shown to be effective in the treatment of amoebiasis, side effects and emergence of parasites resistance prompted the search for novel drug to control this disease. In this regard, the Medicines for Malaria Venture (MMV) Pathogen Box library of selected compounds was screened to identify anti-*Entamoeba histolytica* agents using the resazurin based fluorescence assay. Overall, the results revealed three novel anti-*Entamoeba histolytica* scaffolds with low micromolar activity including MMV675968 (IC_50_ = 2.10 µM), MMV688179 (IC_50_ = 2.38 µM) and MMV688844 (IC_50_ = 5.63 µM). Structure-Activity-Relationship (SAR) studies led to identification of two analogs ∼100 fold more potent and selective than the original hit compound **1** (MMV675968): **1k** (IC_50_ = 0.043 µM) and **1l** (IC_50_ = 0.055 µM). Predictive analysis using Maestro 11.6 suggested that these hit compounds possess acceptable physicochemical and metabolism properties. These lead compounds are therefore good starting points for lead optimization studies towards identification of drug candidate against amoebiasis.

**Author Summary:** Diarrhoea is a leading cause of death for millions of children worldwide. One of the top 15 causes of severe diarrhoea is *Entamoeba histolytica*, causing amoebiasis. What makes *E. histolytica* dangerous is its ability to disseminate easily through a given population via contaminated food and water supplies. Moreover, *E. histolytica* is quite comfortable in the environment, difficult to kill with chorine and infect people at a very low dose, making it a priority pathogen to eradicate. Many drugs have been developed so far to cure this infection. However, they are not efficient enough to control the disease due to pathogen resistance that is becoming a big issue. In addition to that, almost all the drugs in use are highly toxic to human causing several side effects upon medications. Therefore, new, more efficient and less toxic drugs are urgently needed for the better management of amoebiasis. Since the development of a new drug takes years, repurposing existing drugs has been shown to shortcut the process and boost the discovery rate of new medicines. Using this same approach, we have identified two compounds that potently inhibit *E. histolytica* and are nontoxic that can enter the drug discovery pipeline for new amoebicidal drug development. Moreover, these new inhibitors could also serve as starting points for the synthesis of a library of amoebicidal compounds.

## Introduction

Diarrhoea that is credited to have caused approximately 8% of all deaths among children under age 5 worldwide in 2016 is considered as a leading executioner of children. Basically, the number of yearly deaths among children is about 480,000, meaning over 1,300 children dying each day [1]. Amoebiasis caused by the protozoan parasite *Entamoeba histolytica* is listed among the top 15 causes of severe diarrhoea in the first 2 years of life in children living in the developing world [2]. Moreover, approximately 4 to 10% of the carriers of this amoeba infection develop clinical symptoms within a year and amoebic dysentery is considered as the third leading cause of death from parasitic disease worldwide after malaria and schistosomiasis [3-4]. Acute amoebiasis presents symptoms such as diarrhea with frequent and often bloody stools, whereas chronic amoebiasis can present gastrointestinal symptoms plus fatigue, weight loss and occasional fever. Extra-intestinal amoebiasis can occur if the parasite spreads to other organs, most commonly the liver, where it causes amoebic liver abscesses with fever and right upper quadrant abdominal pain [5-6].

Estimates indicate that *E. histolytica* infects approximately 500 million people worldwide, resulting in 50 million cases of invasive disease and about 70,000 deaths annually [7-8]. Because of its easy dissemination through contaminated food and water supplies, in addition to its low infectious dose, chlorine resistance, and environmental stability, *E. histolytica* is classified as a category B priority biodefense pathogen by the National Institute of Allergy and Infectious Diseases [9]. For the control of this infection, the treatment relies on structurally diverse drugs (fig 1) and the administration of the treatment depends on the diagnosis and severity of the illness. Usually, the drugs are effective for the treatment of invasive amoebiasis, but are less effective in eliminating parasites located in the intestinal lumen. Overall, in symptomatic patients and in invasive disease, the most widely used drugs against *E. histolytica* are the nitroimidazoles (metronidazole and tinidazole) [10-12]. However, several side effects are reported, ranging from vomiting and diarrhea, hallucinations [13], encephalopathy [14] to cancer [15] besides emergence of resistant strains that have been reported against these drugs [16]. Therefore, to face these shortcomings, new and better drugs are required to control amoebiasis.

**Fig 1.**
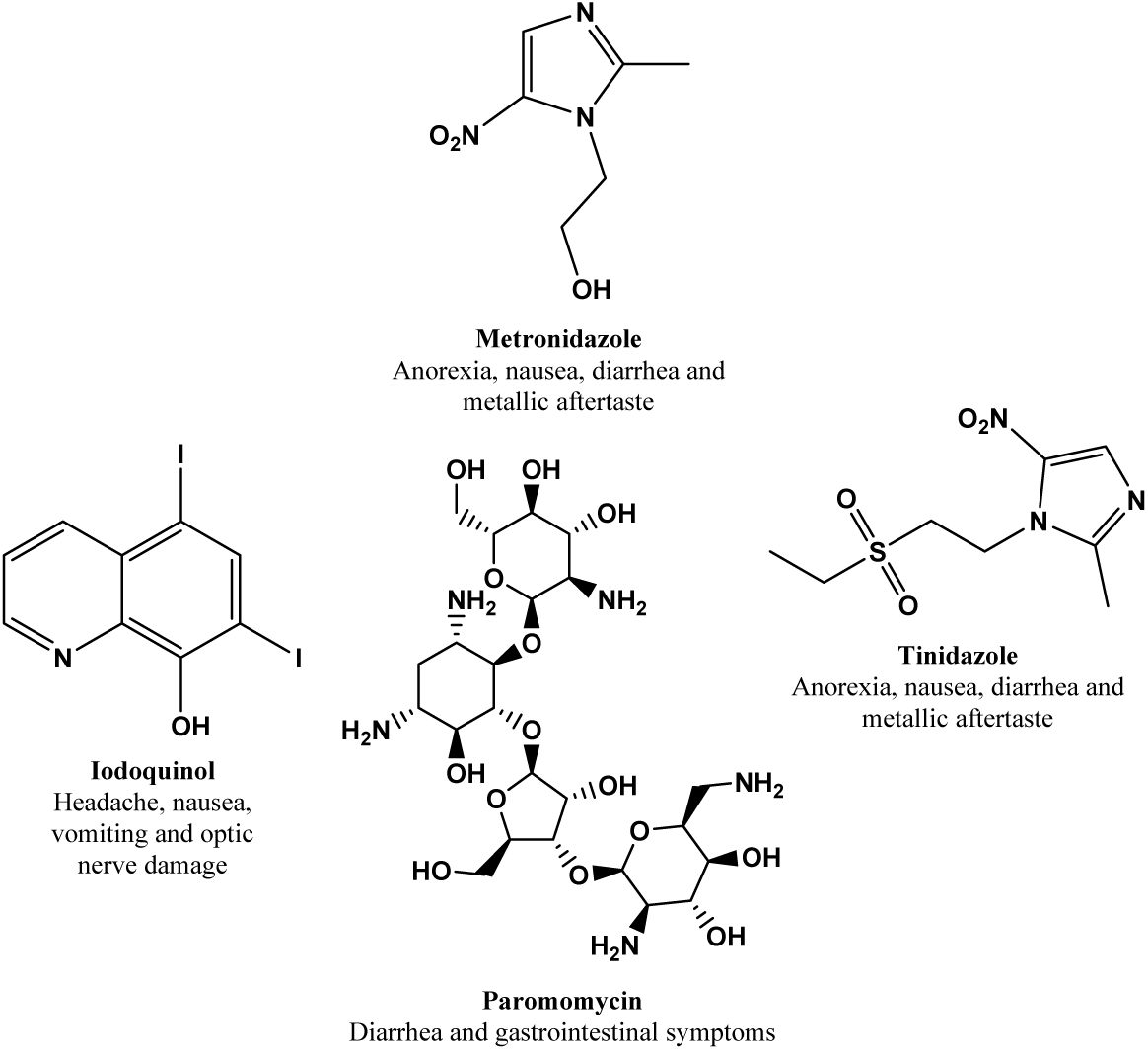
Structure of available drugs against amoebiasis and their respective side effects.

Prompt treatment accounts among the global strategy to control infectious diseases including amoebiasis. However, the efficiency of this intervention highly relies on drugs which should not only be safe and efficacious, but also affordable, and available to the population in need. Ideally, any new anti-amoeba agent should bear all the above characteristics. In the quest for such new treatments against amoebiasis, many strategies have been adopted, including rational drugs design [17] and screening of synthetic and natural products libraries [8, 18]. Several synthesized and naturally occurring compounds have been tested, yet there is no molecule considered to be ideal for the treatment of amoebiasis, particularly for the treatment of severe infections [19]. However, Chacín-Bonilla et al. [20] reported the potential of nitazoxanide prescribed as medication against *Cryptosporidium parvum* and *Giardia lamblia* infection as a key compound for the therapy against both luminal and invasive *E. histolytica* forms, although no further study has been reported since. On another hand, repurposing previously developed drugs or those under development stages against other parasitic or non-parasitic diseases represents a shortest and cheapest way to accelerate the discovery of drugs for neglected tropical diseases, including amoebiasis.

Within this framework, the Medicines for Malaria Venture (MMV) has made available open access libraries of hits generated through screening corporates and academic libraries [21]. The success of this approach went beyond malaria parasites [22-24], where inhibitors of several pathogens of other diseases including tuberculosis [25-26], schistosomiasis [27-29], those caused by kinetoplastids [30], cryptosporidiosis [31], toxoplasmosis [32] and cancers [33] were identified. Based on the same concept, we investigated the MMV Malaria Box which led to identification of compounds exhibiting moderate inhibition against *Entamoeba histolytica* [32]. Following these previous footsteps, we hypothesized that the investigation of the MMV Pathogen Box (MMVPB), consisting in 400 drug-like compounds with low toxicity for mammalian cells and activity against specific microbial pathogens could lead to identification of hit scaffolds different from those of the mainstay treatment to feed and accelerate the discovery and development of new anti-amoebiasis drugs with different modes of action.

## Methods

### Parasites culture and maintenance

The *E. histolytica* strain NR-176 provided by BEI Resources (www.beiresources.org) was maintained in culture in Eagle’s minimum essential medium (EMEM) (Sigma, Germany) supplemented with 10% albumin bovine serum (ABS) and 1% penicillin-streptomycin solution. Parasites were sub-cultured twice weekly. For the assays, cells were harvested by chilling the tube on ice for 15 min to detach the parasites, and then centrifuged at 300×g for 5 min. The supernatant was decanted, and cells pellets were resuspended in fresh medium. The number of viable cells was calculated using a haemocytometer and 0.4% (w/v) trypan blue dye. The criteria for viability were motility and dye exclusion.

### The MMVPB compounds library

The Pathogen Box was kindly provided by Medicines for Malaria Venture (MMV, Switzerland) and consisted in 400 drug-like compounds. Compounds were supplied in 96-well microtiter plate format containing 20µL/well of 10 mM solution dissolved in 100% DMSO. Intermediary solution for each compound was prepared in 96-well plate at 100µM by diluting compounds in incomplete culture medium. All plates were stored at −20°C.

### Assessment of amoebicidal activity of compounds via resazurin reduction assay

#### Determination of *E. histolytica* inoculum size using fluorescence intensity

*E. histolytica* trophozoites at 9.69×10^3^cell/mL, 1.94×10^4^cell/mL, 3.88×10^4^cell/mL, 7.75×10^4^cell/mL, and 1.55×10^5^cell/mL in 100μL were added in triplicate in the 96-wells flat-bottomed plates and incubated for 48 h at 37°C in an atmosphere of 5% CO_2_. Upon 48 h of incubation, 10 μL of resazurin (0.15mg/mL) were added and mixed gently and incubated in the dark at 37°C for 30 min. Fluorescence was subsequently measured using Infinite M200 plate reader (Tecan) with excitation and emission at 530 and 590 nm respectively.

#### Assay validation through Z-factor determination

To validate our resazurin reduction assay, the statistical effect size (Z-factor) was calculated. In a 96 wells plate format, *E. histolytica* (2×10^4^cells/mL) was cultured in Eagle’s minimum essential medium (EMEM) (Sigma, Germany) supplemented with 10% albumin bovine serum (ABS) and 1% penicillin-streptomycin solution for 48 h at 37°C, 5% CO_2_. After 48h incubation, 10 μL resazurin at 0.01% (wt/vol) were added in each well and incubated in the dark for 30 min at 37°C. Plates were scanned using a Tecan Infinite M200 fluorescence multi-well plate reader (Männedorf, Switzerland) with excitation and emission at 530 and 590 nm respectively. More than 30 replicate of the negative control (Parasite-free inhibitor), positive control (1mg/mL metronidazole in 100% DMSO) and the blank (medium-free parasite) were prepared. The assay was performed twice and the data were used to calculate the Z-factor using the formula:

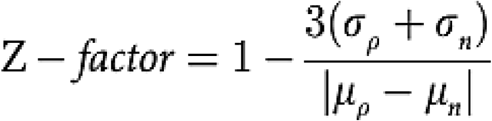

Where *σp* and *σn* are the standard deviations of the positive and negative controls respectively, and *μp* and *μn* are the corresponding mean values. A Z-factor between 0.5 and 1.0 indicates an excellent assay and statistically reliable separation between the positive and negative controls.

#### Determination of Single Point Growth Inhibition of the MMVPB compounds

Screening of the 400 compounds was conducted in 96-well sterile polystyrene flat-bottom microtiter plates (Corning). *E. histolytica* culture at 2×10^4^ cells/mL was exposed to 10 µM drug in a final volume of 100 µL of EMEM per well. Plates were incubated at 37°C under an atmosphere of 5% CO_2_ for 48h. Metronidazole at 1 mg/mL and 0.4% DMSO were used respectively as positive and negative controls. After 48 h of incubation, 10μL of resazurin (0.15 mg/mL) (Sigma, USA) were added to each well, mixed gently, and incubated in the dark at 37°C for 30 min. Subsequently fluorescence was measured using a Tecan Infinite M200 fluorescence multi-well plate reader (Austria) with excitation and emission at 530 and 590 nm, respectively. Tests were performed in three independent experiments. Compounds with inhibition percentage ≥70% were selected for dose-response assay.

#### Dose-Response Growth Inhibition assay of the selected MMVPB Compounds

Median inhibitory concentrations (IC_50_) of the selected MMVPB compounds were determined as described above, with little modifications consisting in tested concentrations ranging from 0.02-25µM for compounds and 1.11-584.24 µM for the reference drug (metronidazole). Experiments were performed in triplicate and repeated twice. Dose–response curves were constructed by plotting mean percent inhibition calculated from the fluorescence counts versus the drug concentrations, and activity was expressed as 50% inhibitory concentration (IC_50_) using the IC Estimator-version 1.2 software.

#### Cytotoxicity study of potent anti-*Entamoeba histolytica* compounds

The cytotoxicity of anti-amoebic compounds was assessed using the MTT assay [34], targeting Vero cells line (ATCC^®^ CCL-81™) cultured in complete medium containing 13.5 g/L EMEM (Gibco, Waltham, MA USA), 10% foetal bovine serum (Gibco, Waltham, MA USA), 0.21% sodium bicarbonate (Sigma-Aldrich, Waltham, MA USA) and 50μg/mL gentamicin (Gibco, Waltham, MA USA). Essentially, Vero cells at 10^4^ cells/200μL/well were seeded into 96-well flat-bottomed tissue culture plates (Corning, USA) in complete medium. Fifty µL of serially diluted compounds solutions (concentrations ≤ 50µM) were added after 24 h of seeding then incubated for 48 h in a humidified atmosphere at 37°C and 5% CO_2_. DMSO was added as negative inhibitor at 0.4% (v/v). Twenty µL of a stock solution of MTT (5mg/mL in 1X phosphate buffered saline) were added to each well, gently mixed, and incubated for an additional 4 h. After spinning the plate at 1,500 rpm for 5 min, the supernatant was carefully removed and 100 μL of 100% DMSO (v/v) were added. Formazan formation was measured on a Magelan Infinite M200 fluorescence multi-well plate reader (Tecan) at 570 nm. The 50% cytotoxic concentrations (CC_50_) of compounds were determined by analysis of dose – response curves using GraphPad Prism 7.0. The selectivity Indices (CC_50_ Mammalian cell/IC_50_ *E. histolytica*) were calculated for each compound.

### Statistical analysis

The data was analysed in Microsoft Excel and Prism 7.0 software (GraphPad Software, San Diego, CA). A nonlinear regression sigmoidal dose-response curve fit was applied to dose-response data for both 50% inhibitory concentration and 50% cytotoxic concentration.

## Results

### Choice of *E. histolytica* inoculum size for the assays

The correlation coefficient of the line was 0.953 (fig 2), indicating a linear response between cell/parasite number and fluorescence values at 530 nm (excitation) and 590 nm (emission).

**Fig 2.**
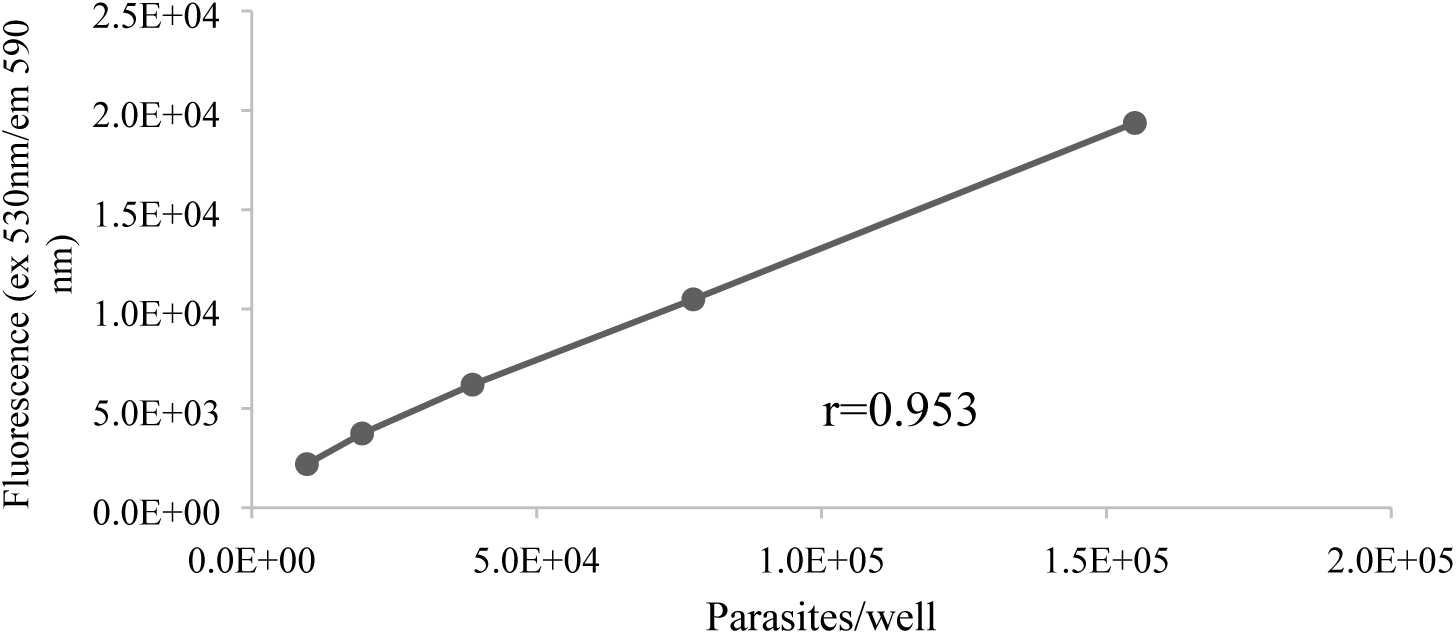
Correlation of *E. histolytica* inoculum size with fluorescence values at 530 nm (excitation) and 590 nm (emission) measured using the Resazurin Assay. Different inoculum sizes of *Entamoeba histolytica* parasites were added in triplicate to the wells of a 96-well plate in EMEM supplemented with 10% ABS, and 1% penicillin-streptomycin. The medium was allowed to equilibrate for 48 h; then 10µl/well of Resazurin Reagent was added. After 30 min at 37°C in a humidified, 5% CO2 atmosphere, the fluorescence at 530 nm (excitation) and 590 nm (emission) was recorded using an Infinite M200 plate reader (Tecan). Each point represents the mean ± SD of 3 replicates.

### Z-factor value for assay validation

The quality of the screen was evaluated using the Z-factor based on the percent inhibition against *E. histolytica* between the 1.0% DMSO and 1mg/mL metronidazole-treated parasites taken as the negative and positive controls respectively. Fig 3 shows the scatter-plot distribution of the percent inhibition for 1.0% DMSO and 1mg/mL metronidazole. The average Z-factor between the 0.4% DMSO and 1mg/mL metronidazole in the 96-well test plates was 0.66 (fig 3) indicating that the assay could reliably separate positive and negative controls. These findings supported the feasibility of our drug screening assay for use in *E. histolytica* screening.

**Fig 3.**
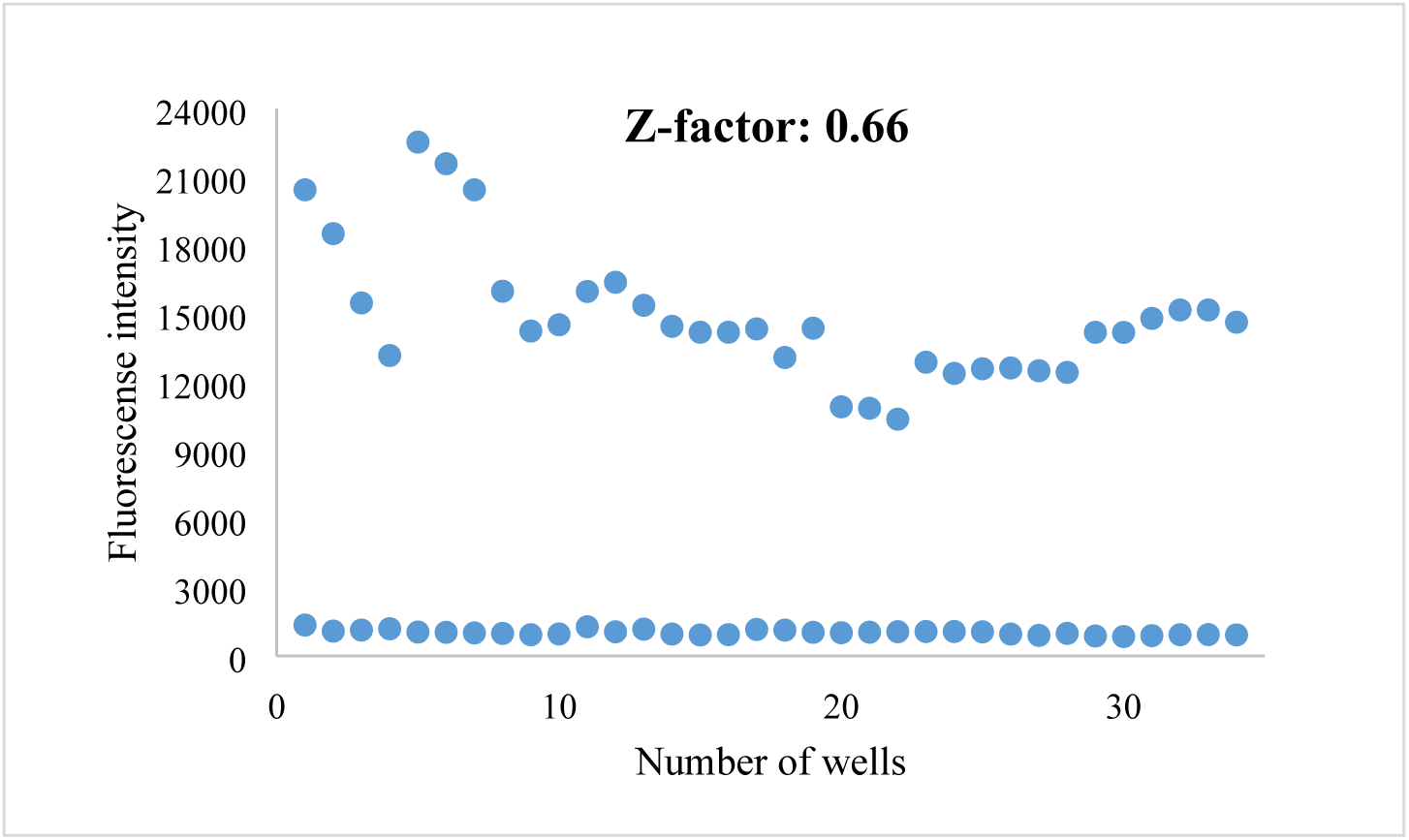
Screening validation. The Z-factor of 0.66 demonstrates an excellent assay respectfully to the guidance defined by Zhang et al. (1999) indicating that for 0.5 ≤ Z < 1, there is a good separation of the distributions between the signal of negative and positive controls, indicating an excellent assay.

### Screening of the Pathogen Box against *E. histolytica* identifies MMV675968, MMV688179 and MMV688844 as potent hits

The preliminary screening of the MMVPB led to the identification of six compounds exhibiting percent inhibition ranging 96-100% against *E. histolytica* in culture including, MMV688978 (Auranofin), MMV688775 (Rifampicin), MMV687798 (Levofloxacin (-)-ofloxacin), MMV675968 (compound **1**), MMV688179 (compound **2**) and MMV688844 (compound **3**). Besides, three other compounds including Linezolid (MMV687803), MMV272144 (compound **4**) and MMV393995 (compound **5**) weakly inhibited the growth of the parasites at the same concentration. The 9 compounds were selected and submitted to dose-response studies as described above. The results achieved indicated, with the exception of compounds **4** and **5**, that 7 compounds could inhibit the growth of *E. histolytica* with IC_50_ values ranging below 10 µM (Table 1).

**Table 1:** Anti-*E. histolytica* activity profile of nine selected MMVPB compounds.

| *MMV ID | Common<br>name or<br>manuscript<br>ID | IC<sub>50</sub> (μM) | #CC<sub>50</sub> (μM) | SI<br>(CC<sub>50</sub>/IC<sub>50</sub>) | Molecular weight<br>(g/mol) | *cLogP | ** Known biological activity |
| --- | --- | --- | --- | --- | --- | --- | --- |
| MMV688978 | Auranofin | 0.07±0.01 | >50 | >714.29 | 678.48 | 1.40 | Rheumatoid arthritis/Amoebiasis |
| MMV688775 | Rifampicin | 0.05±0.00 | >50 | >1000 | 822.94 | 1.29 | Tuberculosis/ Buruli ulcer |
| MMV687798 | Levofloxacin(-<br>)-ofloxacin | 0.045±0.00 | >50 | >1111.11 | 361.37 | -0.34 | Broad-spectrum antibiotic |
| MMV675968 | 1 | 2.10±0.37 | >50 | >23.80 | 359.81 | 2.31 | Cryptosporidiosis |
| MMV688179 | 2 | 2.38±0.64 | 35.43±1.17 | 14.88 | 476.19 | 2.87 | Kinetoplastids |
| MMV688844 | 3 | 5.63±0.6 | >50 | >8.88 | 424.92 | 3.83 | Tuberculosis |
| MMV687803 | Linezolid | 9.17±1.91 | >50 | >5.45 | 337.35 | 0.52 | Tuberculosis |
| MMV272144 | 4 | 13.37±0.68 | 16.98±1.23 | 1.27 | 254.27 | -0.27 | Tuberculosis |
| MMV393995 | 5 | 11.56±0.07 | >50 | >4.32 | 204.23 | 1.10 | Tuberculosis |
| Metronidazole | - | 9.34±0.21 | - | - | 171 | - | Amoebiasis |

\*cLogP= $\log (C_{\text{octanol}}/C_{\text{water}})$ , is the measure of the compound's hydrophilicity; \*\*Known activity of compounds against other diseases. IC<sub>50</sub> obtained from two independent experiments performed in triplicate each; #CC<sub>50</sub> values of compounds against Vero cell line.

Interestingly, among the six compounds that significantly inhibited *E. histolytica* with IC_50_ values ranging 0.045-5.63 µM, three are approved drugs. In fact in this study, Auranofin (MMV688978) completely prevented *E. histolytica* proliferation (100% inhibition) over the incubation time frame. This finding was not surprising given that Auranofin which is used for the management of rheumatoid arthritis has recently been repurposed as drug for the treatment of amoebiasis [18]. As a result, the activity of Auranofin against *E. histolytica* growth further validated our screening approach. Rifampicin (MMV688775) is an important antibiotic drug used for the treatment of buruli ulcer and tuberculosis, and Levofloxacin (-)-ofloxacin (MMV687798), a broad-spectrum antibiotic of the fluoroquinolone class known to exhibit bactericidal activity.

Of particular interest, this study has allowed to identify three hits showing selectivity indexes >8.88 against Vero cell lines and low micromolar activity against *E. histolytica*: IC_50_= 2.10 µM, 2.38 µM, and 5.63 µM for compounds 1, 2 and 3 respectively. These compounds are being reported here for the first time as having amoebicidal activity (Fig 4; Table 1). Given their lipophilicity in an acceptable range (cLogP 2.31-3.83) and small molecular weight (MW 359-476), they might represent novel and attractive anti-*E. histolytica* chemical starting points for medicinal chemistry efforts and therefore deserve further attention.

**Fig 4.**
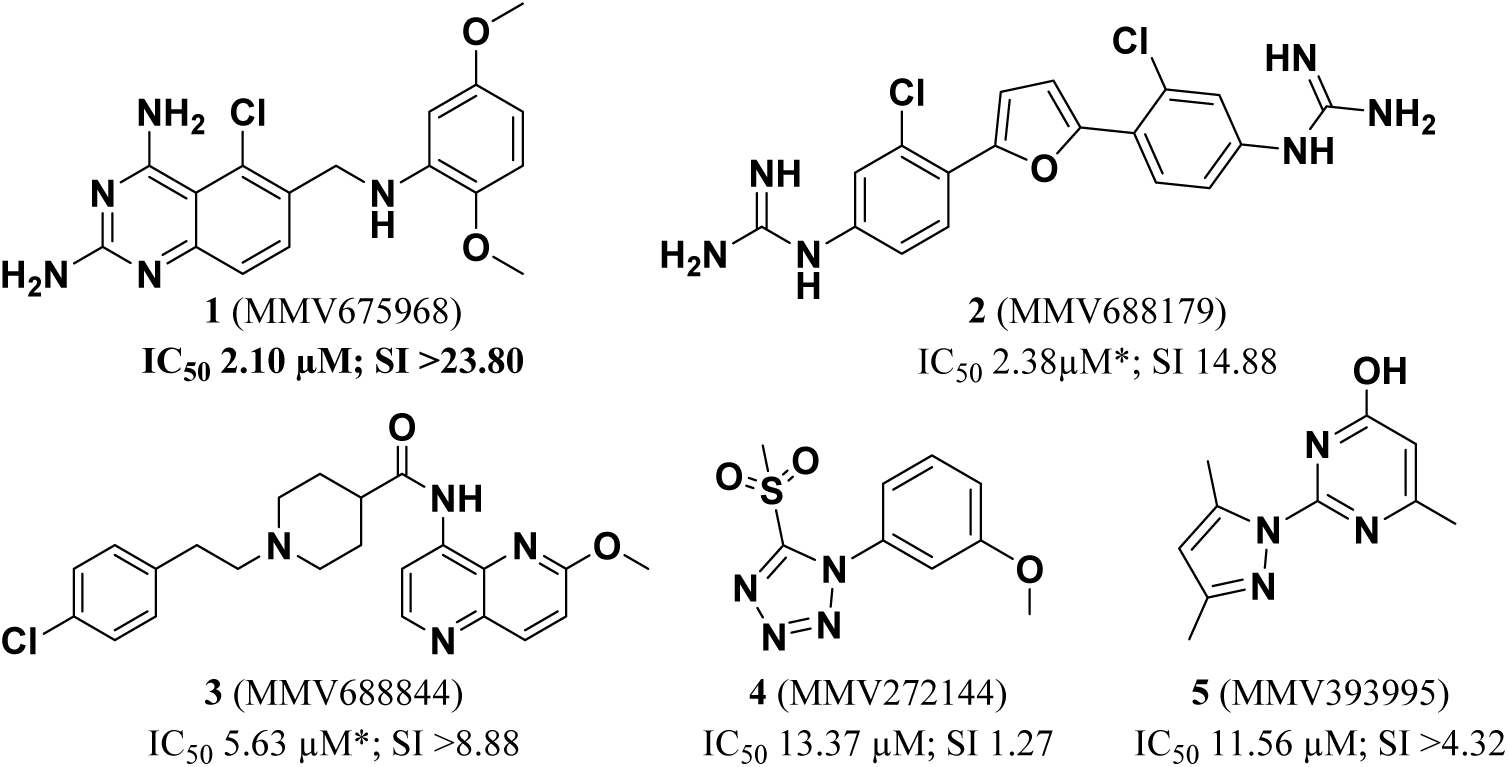
Structures and activity of novel amoebicidal compounds identified from the screening of the Pathogen Box (*activity not reconfirmed on resynthesized hits)

### Screening of resynthesized hits and their analogs identifies two highly potent *E. histolytica* inhibitors: MMV1578523 (1k) and MMV1578540 (1l)

In an attempt to confirm their potency upon retesting, and identify more potent and selective anti-amoebic hits, compounds **1, 2** and **3** were resynthesized and fresh DMSO solutions were prepared from solid samples before testing against *E. histolytica* as described above.

Surprisingly upon testing, the activity was not confirmed for compounds **2** and **3** since a complete loss of inhibitory effect against *E. histolytica* was observed at concentrations below 25µM. Further investigation through available structural analogs of these 2 hits did not show any inhibition against *E. histolytica* while their cytotoxicity values were significant (see Supporting Information). These results suggest that primary activity observed for compounds **2** and **3** might be due to their cytotoxicity or to degradation of the compound under the storage condition of the Pathogen Box. Therefore, they may not be suitable inhibitors against the target parasite. The loss of potency of resynthesized hits is one of the main challenges encountered during hit identification and lead discovery phase of the drug discovery process. This observation led Hughes et al. [35] to define a true ‘hit’ molecule as a compound which has the desired activity in a compound screen and whose activity is confirmed upon retesting. This is as true as the success to identify candidate molecules for clinical development depends upon the stability of the potency over time.

Upon retest, compound 1 showed a 3.6-fold decrease in potency (IC_50_ from 2.100 to 7.495 µM) as well as the selectivity against Vero cell line (SI 5.5). Although slightly disappointed by this loss of potency, this activity level confirmed to be below 10 µM prompted us to test structural analogs of compound **1**. This has allowed to establish rudimentary structure-activity-relationship (SAR) and to identify two analogs ∼100 fold more potent than the parent hit compound.

### Structure activity relationship

Twenty-three analogs of compound **1** were available as solids from MMV as part of the Pathogen Box initiative and tested against *E. histolytica* as an opportunistic approach (see Supporting Information for data on all compounds). The key results allowing to establish very preliminary SAR around compound **1** are summarized in Table 2 below.

**Table 2:**
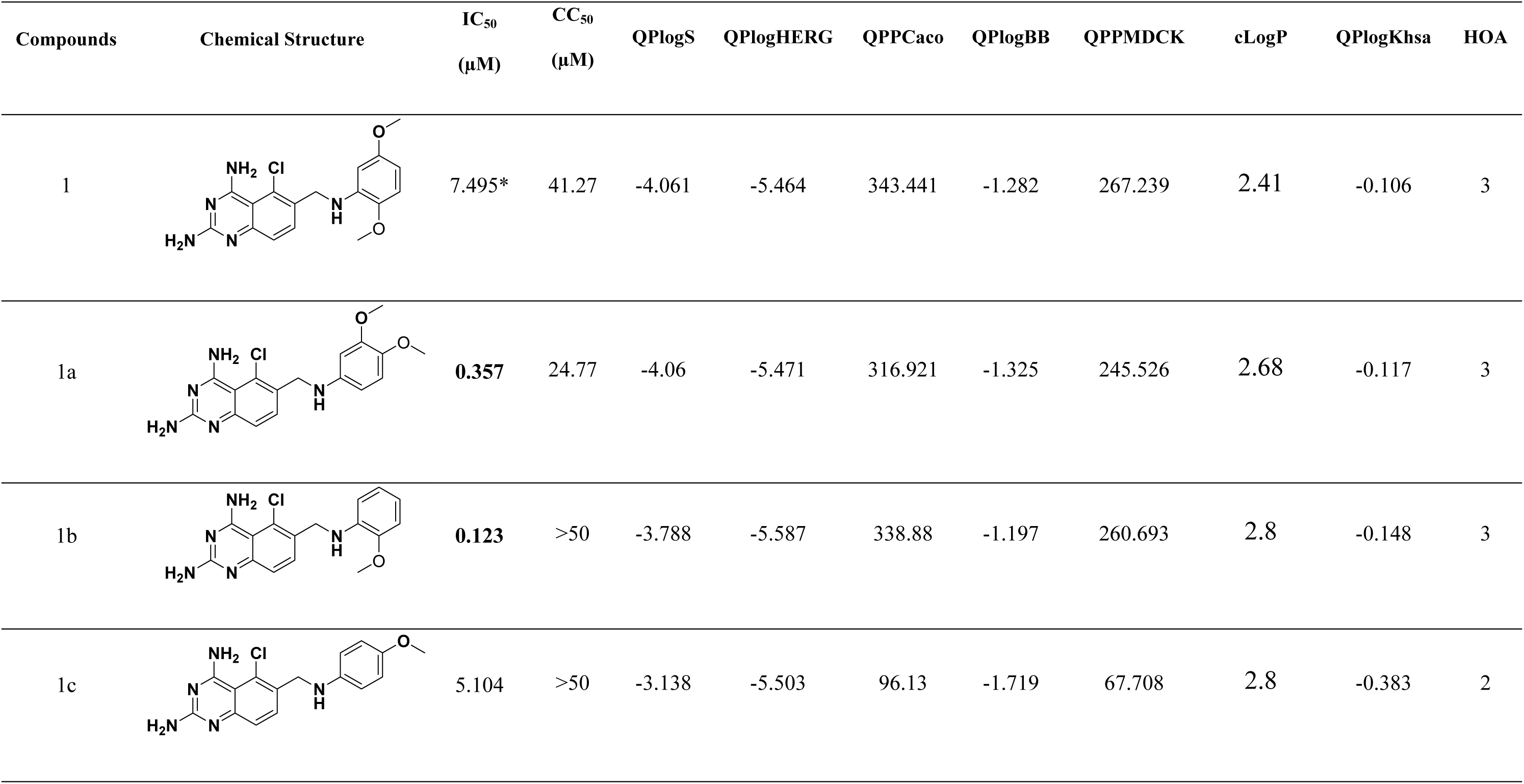
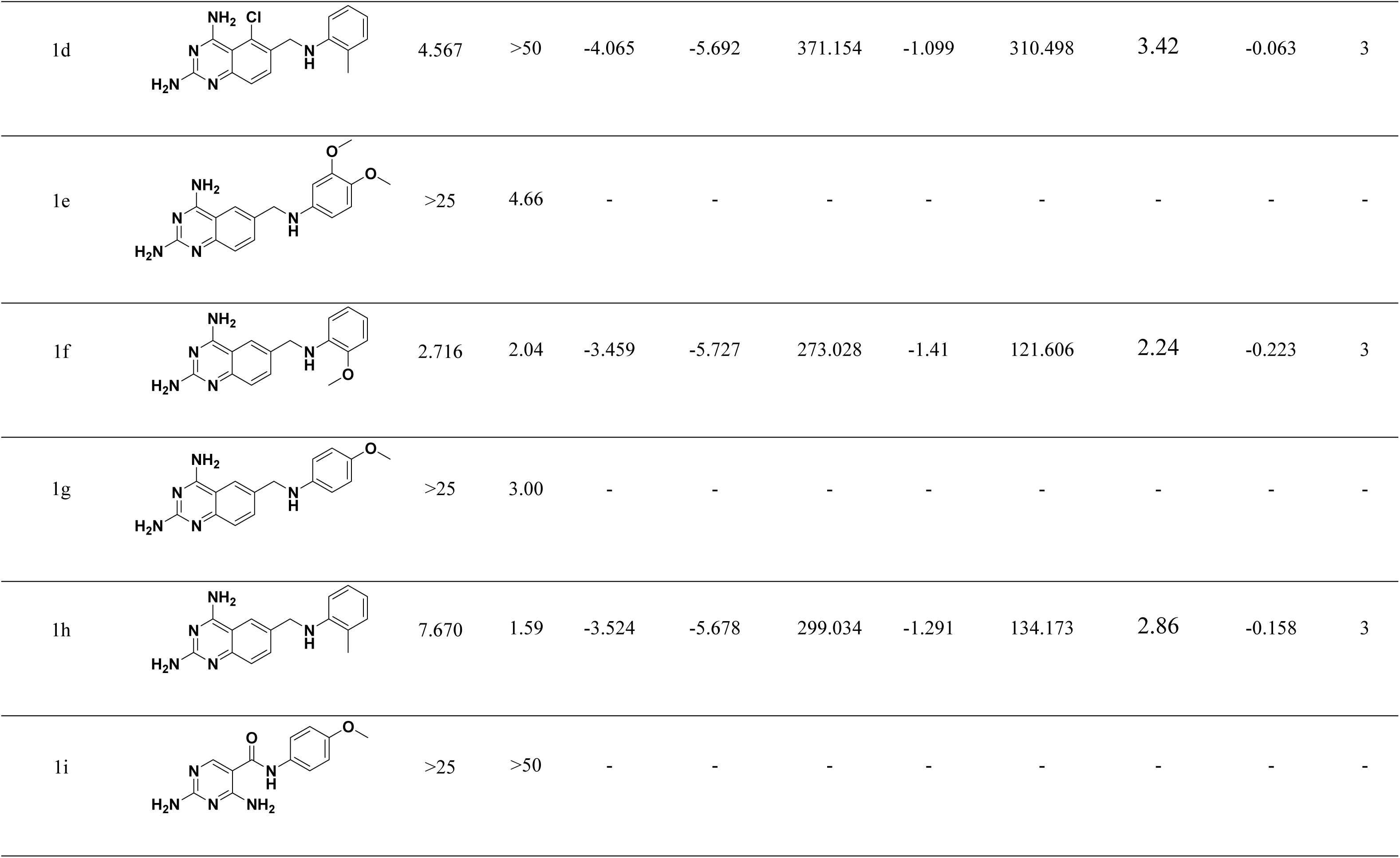
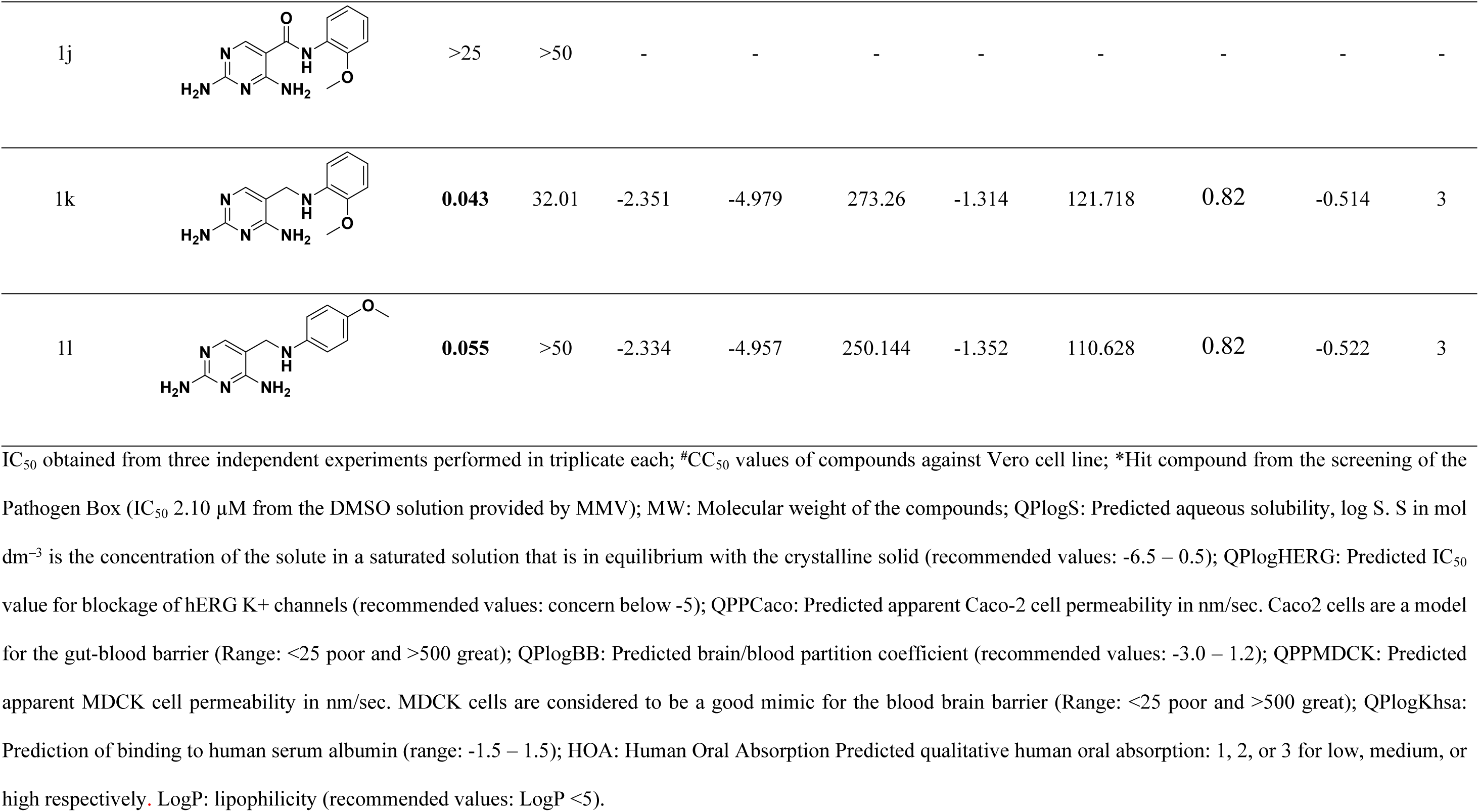
Amoebicidal activity, selectivity and predictive ADME parameters of compound 1 (MMV675968) and structural analogs

Compounds **1a-1d** show different substitution pattern around the phenyl moiety of the hit compound **1** while keeping unchanged the 5-chloro-2,4-diaminoquinazoline core. Among these structural analogs, **1b** was found to be the most potent (IC_50_= 0.123 µM). This 61-fold increase in potency *vs*. the hit compound **1** results from incorporation of an *ortho*-methoxy as the only substituent of the phenyl moiety. Replacement of this methoxy with a methyl group did not result in significant activity improvement (compound **1d**).

The additional analogs depicted in Table 2 deal with modifications of the core moiety. Removal of the chlorine atom from compounds **1a** and **1c** led to complete loss of activity in compounds **1e** and **1g** at concentrations below 25 µM. Moreover, removal of the chlorine atom from the most potent analogue **1b** led to compound **1f**, showing about 22-fold decrease in activity. Overall, these activity changes indicate that the chlorine atom is critical for activity maintenance.

Interestingly, 2,4-diaminopyrimidine as core moiety led to a dramatic improvement in potency with compounds **1k** and **1l** showing 3- and 93-fold activity increase respectively when compared to their 5-chloro-2,4-diaminoquinazoline analogs **1b** and **1c**. Overall, screening of the available analogs of the hit compound **1** has allowed to improve the *in vitro* potency against *E. histolytica* by 147-fold.

**1l** and **1k** constitute promising starting points. However, it would be key to assess the scope for modifications around the phenyl moiety and to determine which functionalities could be tolerated. Removal of the benzylic position in compounds **1k** and **1l** by incorporation of an amide functionality resulted in complete loss of activity for **1i** and **1j**. The length and rigidity of the spacer connecting the diaminopyrimidine core to the methoxyphenyl group deserves also further investigation.

## Discussion

In the quest for new drugs against infectious diseases, repurposing existing active principles is among the novel and fast-track approaches in drug discovery research. This approach significantly reduces the time frame, cost, effort and clinical failure risk associated with conventional drug discovery approaches [36]. In fact, repurposing unsuccessful drug candidates and existing drugs has successfully discover new active approved drugs for other indications [37].

In the present study, some MMVPB compounds have shown highly potent activity against *E. histolytica*, the causative agent of amoebiasis. Auranofin (MMV688978) was identified to exhibit very good amoebicidal activity (IC_50_= 0.07 μM). This result corroborates a recent report of potent activity of Auranofin against *E. histolytica* with IC_50_ value 10-fold better *in vitro* than metronidazole (0.5 vs 5μM) [18], the best treatment for amoebiasis, together with tinidazole.

Auranofin exerts its action through the inhibition of reduction/oxidation (redox) enzymes that are essential for maintaining intracellular levels of reactive oxygen species. This inhibition leads to cellular oxidative stress and intrinsic apoptosis [38-41]. Through this mechanism of action, Auranofin has been reported as potential drug lead against several diseases including cancer, neurodegenerative disorders, HIV/AIDS, parasitic and bacterial infections [41].

Also, two of the hits that were identified in this study, MMV688775 (Rifampicin) and MMV687798 (Levofloxacin (-)-ofloxacin) exhibited potent amoebicidal activities (IC_50_ of 0.05 μM and 0.045 μM respectively). They are well known for their broad-spectrum antibiotic activity against a variety of bacterial pathogens. However, their amoebicidal properties have not been reported before. Three other MMVPB compounds exerted significant activity against *E. histolytica.* Compound **2** (IC_50_ of 2.38 µM) is thought to be a DNA groove binder as likely mechanism of action, and is active against several bacterial and fungal pathogens [42]. Besides, this compound was among the seven MMV Pathogen Box compounds that exhibited bacteriostatic or bactericidal activity against *Burkholderia pseudomallei*, the causative agent of melioidosis, a disease that requires long-term treatment regimens with no assurance of bacterial clearance [43]. To a lesser extent, compound **3** also exhibited promising activity against *E. histolytica* (IC_50_ 5.63µM). It was originally identified as non-cytotoxic *Mycobacterium tuberculosis* hit in GlaxoSmithKline (GSK) whole cell screens and was predicted, based on *in silico* analyses, to target ABC transporters (Rv0194) in *M. tuberculosis* [26, 44] also involved in the resistance mechanisms of several parasitic protozoa including *E. histolytica* [45]. This compound also showed potency against non-tuberculous mycobacteria (*M. abscessus* and *M. avium*) [46-47] and other intestinal protozoan parasites such *Giardia lamblia* and *Cryptosporidium parvum* [48].

Compound **1** (MMV675968) has previously been reported to have anti-cryptosporidiosis activity and to target the dihydrofolate reductase (DHFR) in *Pneumocystis carinii* and *Toxoplasma gondii* [49-50]. Moreover, the chemical structure of compound **1** show some similarity to piritrexim and trimetrexate, two nonclassical folic acid inhibitors approved for the treatment of *Pneumocystis carinii* infection in AIDS patients [51]. A recent interrogation of the MMV Pathogen Box has identified **1** as the most active of the box compounds against *Toxoplasma gondii* with IC_50_ of 0.02µM and a selectivity index of 275 [52]. This compound was also identified as a dual hit of *Cryptosporidium parvum* and *Giardia lamblia* with 88% and 90% inhibition respectively in the initial screen of the box compounds by Hennessey et al. [48]. Another study by Lim et al. [53] has found compound **1** among the 13 most potent MMV Pathogen Box compounds against *Madurella mycetomatis*, a fungus primarily reported in Central Africa as the causative agent of mycetoma in humans, which is a chronic infectious and inflammatory disease. Beyond the numerous activities reported of compound MMV675968, it was also showed in this study to be active against *E. histolytica* with IC_50_ of 2.10µM, rationally inhibiting the amoebal DHFR enzyme as suggested by the findings of Lau et al. [50]. In fact, competitive inhibitors of DHFR are used in the chemotherapy or prophylaxis of many protozoan pathogens, including the eukaryotic parasites *Plasmodium falciparum, Entamoeba histolytica* and *Toxoplasma gondii* [50].

It is likely that lead compounds identified in this study inhibits *Entamoeba histolytica* through inhibition of the enzyme dihydrofolate reductase. In fact, the enzyme dihydrofolate reductase (DHFR) catalyzes the reduction of folate to dihydrofolate(DHF) and DHF to tetrahydrofolate (THF) by use of the cofactor NADPH. The methylenated form of THF serves as a carbon donor for the synthesis of thymidylic acid in a reaction catalyzed by thymidylate synthase (TS). Knowing that thymidylic acid is essential for DNA synthesis, blocking of either DHFR or TS activity will lead to cell death [54]. Indeed, success of DHFR inhibitors in treating various infectious diseases can be attributed to the divergence in the DHFR sequence, which imparts a high degree of selectivity for certain antifolates for one organism versus others. This could be particularly important for parasitic protozoa, which, unlike humans, express DHFR as part of a bifunctional enzyme containing both DHFR and TS activity in two domains of the same polypeptide joined by a linker [55]. This structural and mechanistic distinction of protozoan DHFRs offers a unique opportunity to develop new drug with greater selectivity [51].

Considered as a promising starting point for drug discovery against *E. histolytica*, a preliminary hit expansion was performed in this study through the screening of twenty-three structural analogs of compound **1** to establish rudimentary structural activity relationship. Four analogs displayed good potency, including compounds **1k** and **1l** that were >100-fold more potent than the original hit. DMPK profiling, additional modifications to understand the SAR and to deal with possible metabolic hotspots (e.g. methoxy group, benzylic position) would be required as next steps to assess further the potential of this chemotype.

Overall, all the novel anti-*E. histolytica* hits identified from this study are structurally different from currently available drugs. Furthermore, they are selective against Vero cell lines and have favorable physicochemical properties. Very preliminary structure-activity-relationship has allowed to identify double-digit nanomolar inhibitors against *E. histolytica* in cellular assays. This suggests that medicinal chemistry efforts focusing on lead optimization could result in successful selection of a drug candidate for amoebiasis drug development.

## Acknowledgments

Authors are very grateful to the strong institutional support from the University of Yaoundé 1, Cameroon.

MMV supported this work through a Pathogen Box Challenge grant (PO 15/01083[03]) to Prof. Boyom and provided the open access Pathogen Box and structural analogs.

*Entamoeba histolytica* strain NR-176 was obtained from BEI Resources, NIAID, NIH.

This work also received materials and equipment support from the Yaoundé-Bielefeld Bilateral Graduate School for Natural Products with Antiparasite and Antibacterial Activity (YaBiNaPA).

The study was also supported by the Seeding Labs’ Instrumental Access Grant (SL2012-2) to Prof. Boyom.

## Supporting Information Legends

**Table:** Amoebicidal potency of the hit compounds and their structural analogues.

IC_50_ = Median inhibitory concentration of active compounds against *E. histolytica*; CC_50_ = Median cytotoxic concentration of active compounds against Vero cells; SI (CC_50_/IC_50_) = Selectivity index

